# Therapeutic Potential of Blocking GAPDH Nitrosylation with CGP3466b in Experimental Autoimmune Encephalomyelitis

**DOI:** 10.1101/2022.08.31.505712

**Authors:** Wesley H. Godfrey, Soonmyung Hwang, Payam Gharibani, Efrat Abramson, Michael D. Kornberg

## Abstract

Multiple sclerosis (MS) is a neuroinflammatory disease of the central nervous system (CNS). Although classically considered a demyelinating disease, neuroaxonal injury occurs in both the acute and chronic phases and represents a pathologic substrate of disability not targeted by current therapies. Nitric oxide (NO) generated by CNS macrophages and microglia contributes to neuroaxonal injury in all phases of MS, but candidate therapies that prevent NO-mediated injury have not been identified. Here, we demonstrate that the multifunctional protein glyceraldehyde-3-phosphate dehydrogenase (GAPDH) is robustly nitrosylated in the CNS in the experimental autoimmune encephalomyelitis (EAE) mouse model of MS. GAPDH nitrosylation is blocked in vivo with daily administration of CGP3466b, a CNS-penetrant compound with an established safety profile in humans. Consistent with the known role of nitrosylated GAPDH (SNO-GAPDH) in neuronal cell death, blockade of SNO-GAPDH with CGP3466b attenuates neurologic disability and reduces axonal injury in EAE independent of effects on the immune system. Our findings suggest that SNO-GAPDH contributes to neuroaxonal injury during neuroinflammation and identify CGP3466b as a candidate neuroprotective therapy in MS.

## 1 Introduction

Multiple sclerosis (MS) is an autoimmune disease of the central nervous system (CNS) that affects nearly 1 million people in the United States (Wallin et al., 2019). MS is classically characterized by inflammatory attack against oligodendrocytes, which are the myelin-producing cells crucial for efficient transduction of action potentials along axons, leading to demyelination. The disease is further characterized by gliosis and neurodegeneration (Longo et al., 2018). The two main types of MS are relapsing-remitting MS (RRMS), which is characterized by discrete neurologic “relapses” associated with focal inflammatory attack within the CNS, and progressive MS defined by insidious disability progression in the absence of relapse. Despite a historical focus on demyelination, neuroaxonal loss occurs in all phases of MS and represents a primary source of neurologic disability (Friese et al., 2014), with distinct but overlapping immune mechanisms producing injury in RRMS versus progressive MS (Chitnis & Weiner, 2017; Frischer et al., 2009; Grigoriadis & van Pesch, 2015; Lassmann et al., 2012; Mahad et al., 2015). Currently approved therapies target the peripheral immune system to limit relapses, but none mitigate neuroaxonal injury during relapse or slow the accelerating neurodegeneration associated with progressive MS (Wei et al., 2021). Therefore, a significant unmet clinical need exists for neuroprotective therapies that prevent inflammatory neuroaxonal damage.

During inflammation, a variety of free radicals are generated (Biswas et al., 2017). One particularly potent free radical is nitric oxide (NO), a gaseous, highly reactive molecule associated with inflammatory damage (Sharma et al., 2007). The primary source of NO in neuroinflammation is microglia and CNS-resident macrophages, which contribute to neuroaxonal injury in both relapsing and progressive forms of disease (Friese et al., 2014; Grigoriadis & van Pesch, 2015). NO produced by activated microglia and macrophages plays a key role in MS pathology (Lassmann et al., 2012; Mahad et al., 2015), as it is found in high concentrations within inflammatory MS lesions (Bo et al., 1994; Smith & Lassmann, 2002). In experimental autoimmune encephalomyelitis (EAE), the primary mouse model of MS, locally administered NO scavengers prevent axonal degeneration (Nikić et al., 2011), and NO donors replicate EAE-like axonal pathology (Smith et al., 2001). However, no current therapies target NO-mediated CNS injury.

A primary mode of action of NO signaling is protein S-nitrosylation, a redox-based modification of cysteine residues within target proteins (Hess et al., 2005; Kovacs & Lindermayr, 2013). The glycolytic enzyme glyceraldehyde-3-phosphate dehydrogenase (GAPDH) is physiologically nitrosylated at its Cys150 residue, which inactivates its enzyme activity and produces a role in signal transduction mediated by binding to Siah1 and nuclear translocation (Hara et al., 2005). In the nucleus, nitrosylated GAPDH (SNO-GAPDH) leads to cell death via p300/CBP and p53 pathways (Hara et al., 2005; Sen et al., 2008) and trans-nitrosylates nuclear proteins (Kornberg et al., 2010), with these studies demonstrating a role in neuronal injury. Furthermore, GAPDH nitrosylation was previously reported in the context of neuroinflammation (Bizzozero & Zheng, 2009). We therefore wondered whether blocking GADPH nitrosylation might be neuroprotective in the context of neuroinflammatory disease.

CGP3466b (Omigapil) is a CNS-penetrant compound with broad anti-apoptotic properties that is structurally similar to the anti-parkinsonian drug L-deprenyl (selegiline) but without effects on monoamine oxidase (Harraz & Snyder, 2015; Waldmeier et al., 2000). Notably, CGP3466b blocks GAPDH nitrosylation without affecting GAPDH enzyme activity, and blockade of GAPDH nitrosylation is crucial for the drug’s anti-apoptotic effect (Hara et al., 2006; Kragten et al., 1998). CGP3466b showed exceptional promise at eliciting neuroprotection in pre-clinical mouse models of Parkinson’s disease (PD) and amyotrophic lateral sclerosis (ALS) but unfortunately failed to meet primary endpoints in clinical trials (Miller et al., 2007; Olanow et al., 2006). While nitrosative stress plays a large role in the pre-clinical animal models of these diseases, the factors contributing to the respective human diseases are very complex (Konnova & Swanberg, 2018; Sorenson, 2005). In contrast, MS pathology in humans is largely driven by inflammation, which is a major source of NO, and there is strong evidence for the involvement of NO in axonal damage in human MS (Lassmann et al., 2012; Mahad et al., 2015; Parkinson et al., 1997). As such, there is a strong biological rationale for exploring CGP366b as a neuroprotective therapy in MS.

No animal model fully captures the complex neurological and immunological milieu found in MS (Procaccini et al., 2015). However, in this paper we model key aspects of MS pathology with MOG_35-55_ EAE. This model involves immunizing C57BL/6 mice with a myelin oligodendrocyte glycoprotein (MOG)-derived peptide in combination with complete Freund’s adjuvant and pertussis toxin (Constantinescu et al., 2011), initiating an inflammatory immune response targeting CNS myelin and producing quantifiable neurologic deficits beginning between post-immunization day (PID) 10 and 15. In addition to demyelination, MOG_35-55_ EAE is associated with significant neuroaxonal injury which is dependent on NO (Nikic et al., 2011), making it an ideal model in which to study inflammatory neurodegeneration. Importantly, MOG_35-55_ EAE has proven useful in clinical translation, playing a key role in the identification and development of several currently approved MS therapies (Robinson et al., 2014).

In this study, we explore the potential of CGP3466b as a novel therapy for MS. We show that GAPDH is robustly nitrosylated within the CNS during MOG_35-55_ EAE, which is prevented by systemic CGP3466b administration. We further demonstrate that CGP3466b is directly neuroprotective *in vivo*, ameliorating EAE disease severity and reducing neuroaxonal damage in the optic nerve. Finally, we show that CGP3466b acts independently of the peripheral immune system, having minimal effects on immunophenotype *in vitro* and *in vivo*. We conclude that CGP3466b holds promise as an adjunctive neuroprotective therapy in MS.

## 2 Results

### 2.1 Neuroinflammation leads to GAPDH nitrosylation within the CNS in the MOG_35-55_ EAE mouse model of MS, which is prevented by CGP3466b treatment

To determine whether SNO-GAPDH might play a role in neuroinflammatory injury, we first examined whether GAPDH nitrosylation occurs within the CNS during the course of MOG_35-55_ EAE. Using the well described biotin switch assay for detection of nitrosylated proteins (Jaffrey & Snyder, 2001), we found that GAPDH nitrosylation increased dramatically in spinal cord (the primary site of neuroinflammation in EAE) with a time course that matched the CNS inflammatory response (Figure 1A; Supplementary Figure 1A). SNO-GAPDH levels increased beginning at the onset of neurologic symptoms (PID 10) and reached maximal levels at peak disease (PID 15), representing the peak of neuroinflammation and neurologic disability. Resolution of neuroinflammation (PID 28) was associated with a return of SNO-GAPDH levels to baseline. These results suggest that GAPDH nitrosylation is a major consequence of neuroinflammation in a mouse model of MS.

**Figure 1.**
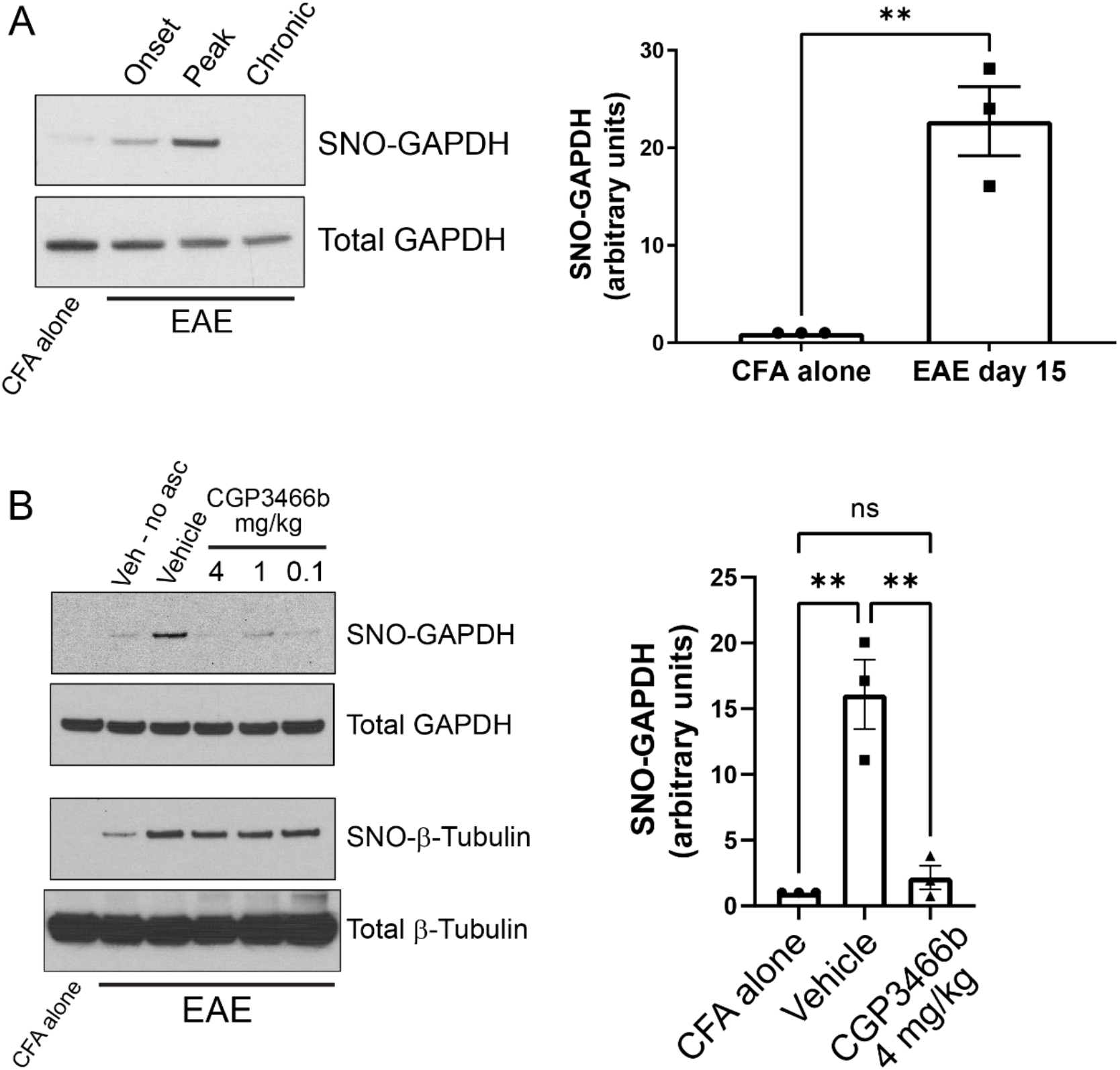
CGP3466b prevents GAPDH nitrosylation within the CNS in the MOG_35-55_ EAE mouse model of MS. **(A)** Nitrosylated GAPDH (SNO-GAPDH) was assayed by biotin switch from spinal cord lysates in control (CFA alone) mice at day 15 and at onset (post-immunization day 10), peak (post-immunization day 15), and chronic (post-immunization day 28) stages of EAE. (*Left*) Representative immunoblot. (*Right*) Quantification of SNO-GAPDH at peak EAE, with SNO-GAPDH normalized to total GAPDH. Data represent mean ± SEM of three mice per group. **(B)** CGP3466b was administered at indicated doses by daily i.p. injection beginning on post-immunization day 0, and nitrosylated GAPDH and β-tubulin were assayed by biotin switch on post-immunization day 15. (*Left*) Representative immunoblot. “No asc” = no ascorbate control. (*Right*) Quantification of SNO-GAPDH levels following treatment with vehicle or CGP3466b 4 mg/kg. Data represent mean ± SEM of three mice per group. ** p<0.01 by student’s t-test (A) or one-way ANOVA with Tukey’s multiple comparisons test (B). Full blots are shown in Supplementary Figure 1.

We then sought to determine whether GAPDH nitrosylation in EAE could be prevented with systemic administration of CGP3466b, similar to other experimental models (Xu et al., 2013; Harraz et al., 2016). Mice were treated with vehicle or CGP3466b daily via intraperitoneal (i.p.) injection beginning on PID 0, and GAPDH nitrosylation was measured by biotin switch assay from spinal cord lysates at PID 15. We found that CGP3466b treatment prevented GAPDH nitrosylation, with maximal effect at 4 mg/kg (Figure 1B; Supplementary Figure 1B). Blockade of nitrosylation was specific to SNO-GAPDH, as CGP3466b had no effect on nitrosylation of β-tubulin.

### 2.2 Blocking SNO-GAPDH with CGP3466b is neuroprotective in the EAE model of multiple sclerosis

Given the known role of SNO-GAPDH in neuronal cell injury (Hara et al., 2005; Sen et al., 2008; Sen et al., 2009; Guha et al., 2016), we examined whether blockade of GAPDH nitrosylation with CGP3466b attenuates neurologic deficits in EAE. To answer this question, we first utilized a prophylactic treatment paradigm, in which mice were randomized to daily treatment with vehicle or CGP3466b beginning on the day of MOG_35-55_ immunization (PID 0). Based on our previous findings, we used the dose of CGP3466b (4 mg/kg) that maximally prevented GAPDH nitrosylation in this model. Mice were clinically scored in a blinded fashion through PID 31. In this treatment paradigm, we found that CGP3466b significantly attenuated the severity of EAE (Figure 2A).

**Figure 2.**
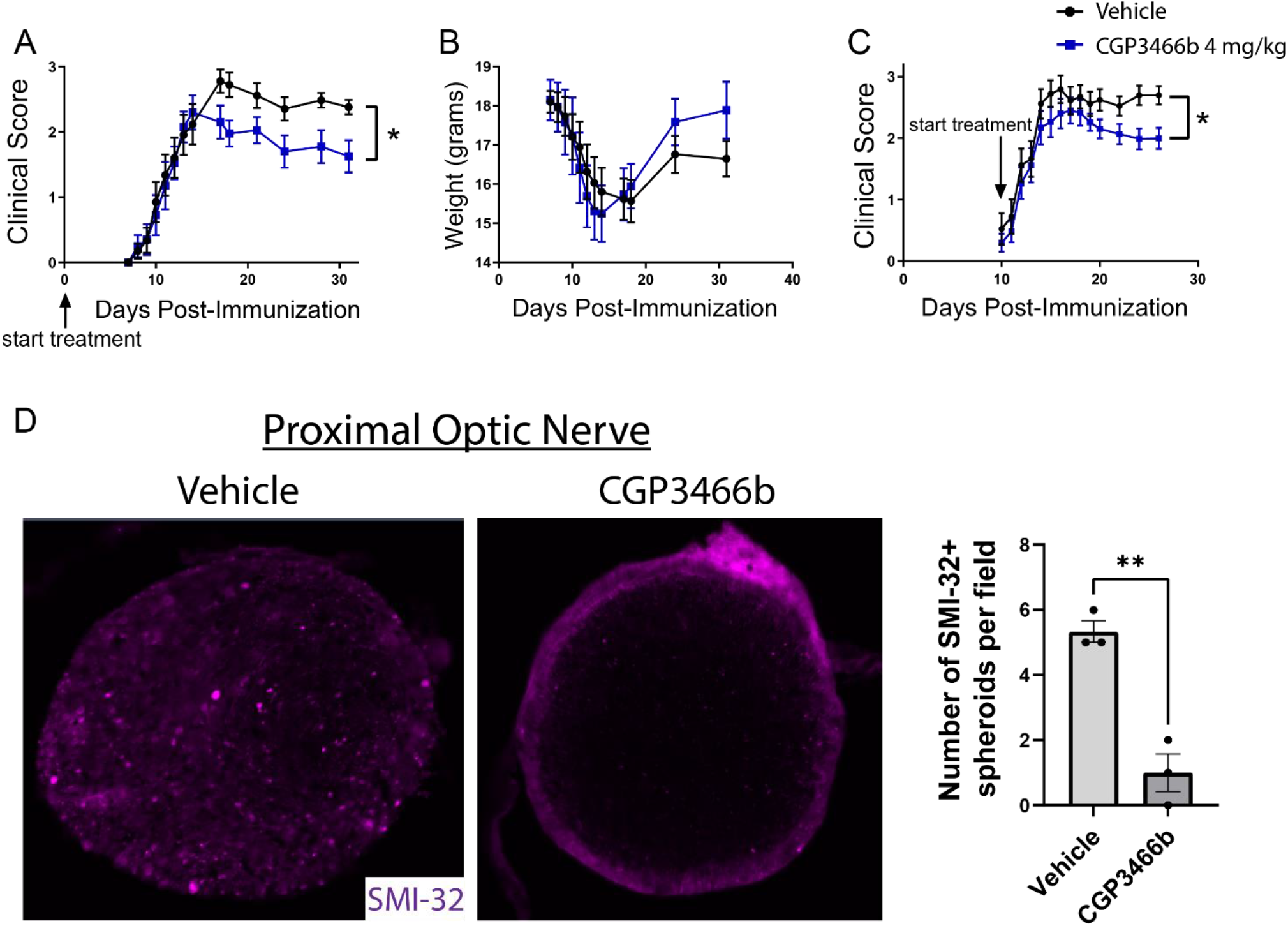
Blocking SNO-GAPDH with CGP3466b is neuroprotective in MOG_35-55_ EAE. **(A and B)** CGP3466b 4 mg/kg was administered by daily i.p. injection beginning on post-immunization day 0 (prophylactic paradigm). Clinical scoring was performed by a blinded observer. Prophylactic treatment with CGP3466b attenuated neurologic deficits (A) without impacting weight loss (B). Data represent n=17 (vehicle) and n=10 (CGP3466b) mice per group. **(C)** CGP3466b 4 mg/kg was administered by daily i.p. injection beginning on post-immunization day 10 (therapeutic paradigm), and clinical scoring was performed by a blinded observer. Data represent n=20 (vehicle) and n=26 (CGP3466b) mice per group. **(D)** To quantify axonal injury, mice were treated with vehicle or CGP3466b 4 mg/kg by daily i.p. injection beginning on post-immunization day 0, and SMI-32+ axonal spheroids were examined via immunofluorescence staining of proximal optic nerve at post-immunization day 28. A representative image is shown (*left*), along with quantification performed from n=3 mice per group (*right*). *p<0.05, **p<0.01 by Mann-Whitney U test (A and C) or student’s t-test (D). Data shown as mean ± SEM.

In MOG_35-55_ EAE, the myelin-directed immune response begins in the peripheral immune system, with dendritic cells presenting MOG-derived peptide to CD4 cells in peripheral lymph nodes and spleen. These peripherally activated CD4 cells then infiltrate the CNS and activate local macrophages and microglia to produce demyelination and neuroaxonal injury. This peripheral immune activation determines the onset of neuroinflammation and produces systemic weight loss that precedes neurologic deficits. In the prophylactic treatment paradigm described above, treatment with CGP3466b had no effect on the timing of disease onset or the degree of weight loss (Figure 2B), suggesting that the attenuated neurologic deficits observed with CGP3466b might be due to direct neuroprotection from inflammatory injury rather than peripheral anti-inflammatory effects. To further explore this possibility, we examined the impact of CGP3466b when treatment began on PID 10, a time point subsequent to priming of the peripheral immune response. This paradigm also more closely approximates a therapeutic clinical scenario, in which treatment begins after the onset of disease. We found that daily treatment with CGP3466b beginning on PID 10 similarly attenuated neurologic disability at PID 28 (Figure 2C).

To determine the impact of SNO-GAPDH blockade with CGP3466b on neuroaxonal integrity directly, we examined the optic nerve. The optic nerve is a primary site of neuroinflammation and inflammatory axonal injury in EAE, in analogy to optic neuritis occurring in humans as a consequence of MS (Kezuka et al., 2011; Sättler et al., 2008). Axonal damage can be readily quantified in the optic nerve during EAE with SMI-32 immunofluorescent staining for non-phosphorylated neurofilament heavy chain, which labels injured axonal spheroids (Gharagozloo et al., 2021). We initiated daily treatment with vehicle or 4 mg/kg CGP3466b on PID 0 in mice subjected to MOG_35-55_ EAE and quantified SMI-32+ axonal spheroids in the proximal optic nerve on PID 28. CGP3466b-treated mice had fewer SMI-32+ axonal spheroids, indicating reduced axonal injury with CGP3466b treatment (Figure 2D).

### 2.3 CGP3466b has minimal direct effects on myeloid and lymphoid cells *in vitro*

Although the above findings suggest that CGP3466b attenuates neurologic disability in EAE through direct neuroprotection rather than peripheral anti-inflammatory effects, we next asked whether CGP3466b has a quantifiable impact on the activation of cultured immune cells. Beginning with bone marrow derived macrophages (BMMs), we found that CGP3466b had no effect on LPS-stimulated secretion of IL-6, IL-1β, or IL-10 over a range of doses, as measured by enzyme-linked immunosorbent assay (ELISA) (Figure 3A). We then evaluated the effect of CGP3466b on LPS-stimulated expression of the activation markers CD86 and MHC II in BMMs (Figure 3B) and bone marrow derived dendritic cells (BMDCs) (Figure 3C) by flow cytometry. CGP3466b had a minimal effect only at the highest dose.

**Figure 3.**
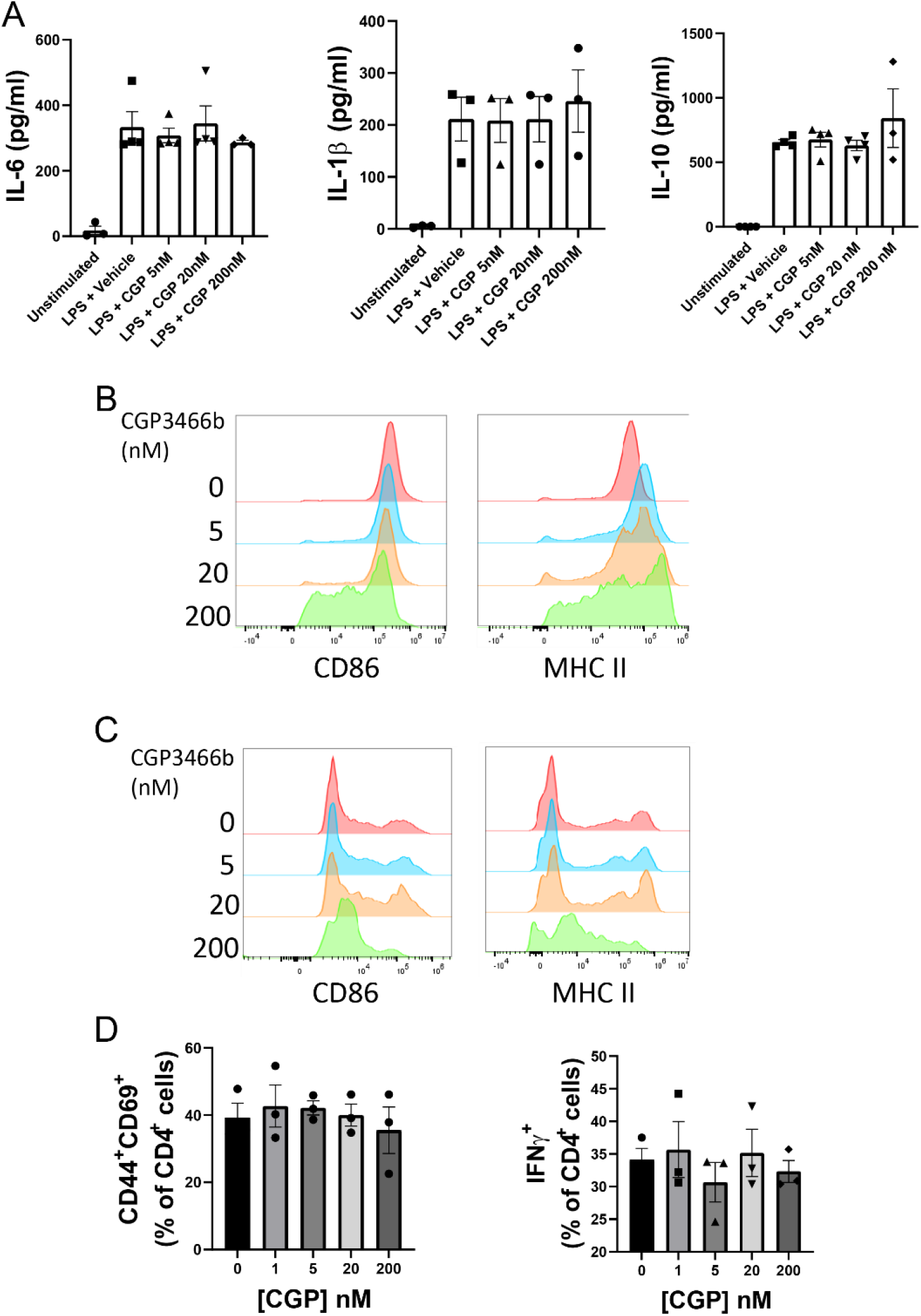
CGP3466b has minimal direct effects on myeloid and lymphoid cells *in vitro*. **(A)** Bone marrow derived macrophages (BMMs) were treated with LPS (100 ng/mL) plus vehicle or the indicated concentrations of CGP3466b for 24 hours. Cytokine concentrations in media were determined by ELISA. Data represent mean ± SEM of n=3 or n=4 independent experiments performed in triplicate. **(B)** BMMs and **(C)** Bone marrow derived dendritic cells (BMDCs) were treated with LPS (100 ng/mL) plus vehicle or the indicated concentrations of CGP3466b, and expression of activation markers was measured by flow cytometry. Data are a representative example of n=3 independent experiments. **(D)** CD4+ lymphocytes were stimulated with anti-CD3/CD28 antibodies (3 ug/mL) in the presence of vehicle or the indicated concentrations of CGP3466b. Percentage of CD4+ lymphocytes expressing the given activation markers was measured by flow cytometry. Data represent mean ± SEM of three biological replicates.

To determine whether CGP3466b directly impacts T cell activation, we stimulated CD4+ T cells with anti-CD3 and CD28 antibodies in the presence of varying concentrations of drug (Figure 3D). CGP3466b had no effect on interferon-gamma (IFNγ) production or expression of the activation markers CD44 and CD69.

### 2.4 CGP3466b does not alter CNS immune infiltration in the MOG_35-55_ EAE model

To further examine whether the neuroprotection provided by CGP3466b occurs independent of peripheral anti-inflammatory effects, we quantified the impact of CGP3466b treatment on CNS inflammatory infiltrates at the peak of EAE. Mice subjected to MOG_35-55_ EAE were treated daily with vehicle or CGP3466b (4 mg/kg) in a prophylactic paradigm (beginning PID 0) and analyses were performed at PID 18. Using immunofluorescence staining of CD45 (expressed by all classes of leukocyte), we detected no difference in the number of infiltrating immune cells in the spinal cord with CGP3466b treatment (Figure 4A). We also examined the impact of CGP3466b on myeloid and lymphoid cell phenotypes in the CNS using flow cytometry. CGP3466b treatment had no effect on arginase-1 (Arg1) expression in either infiltrating macrophages (CD11b^+^CD45^+^Clec12a^+^) (Figure 4B) or microglia (CD11b^+^CD45^+^Clec12a^-^) (Figure 4C) within the brain or spinal cord. iNOS, the classical pro-inflammatory macrophage/microglia phenotypic marker, is only expressed at the onset of EAE (Giles et al., 2018). Consequently, we examined the pro-inflammatory phenotype by examining the effect of CGP3466b on MHC II expression in macrophages (Figure 4D) and microglia (Figure 4E) and found no significant effect. Similarly, we found no significant difference in the proportion of CD4 lymphocytes displaying markers of T helper (Th) 1 (CD3^+^CD4^+^IFNy^+^) (Figure 4F) or Th17 (CD3^+^CD4^+^IL-17^+^) phenotypes (Figure 4G). Together, these results indicate that CGP3466b has minimal effects on the immune response in the MOG_35-55_ EAE model of MS.

**Figure 4.**
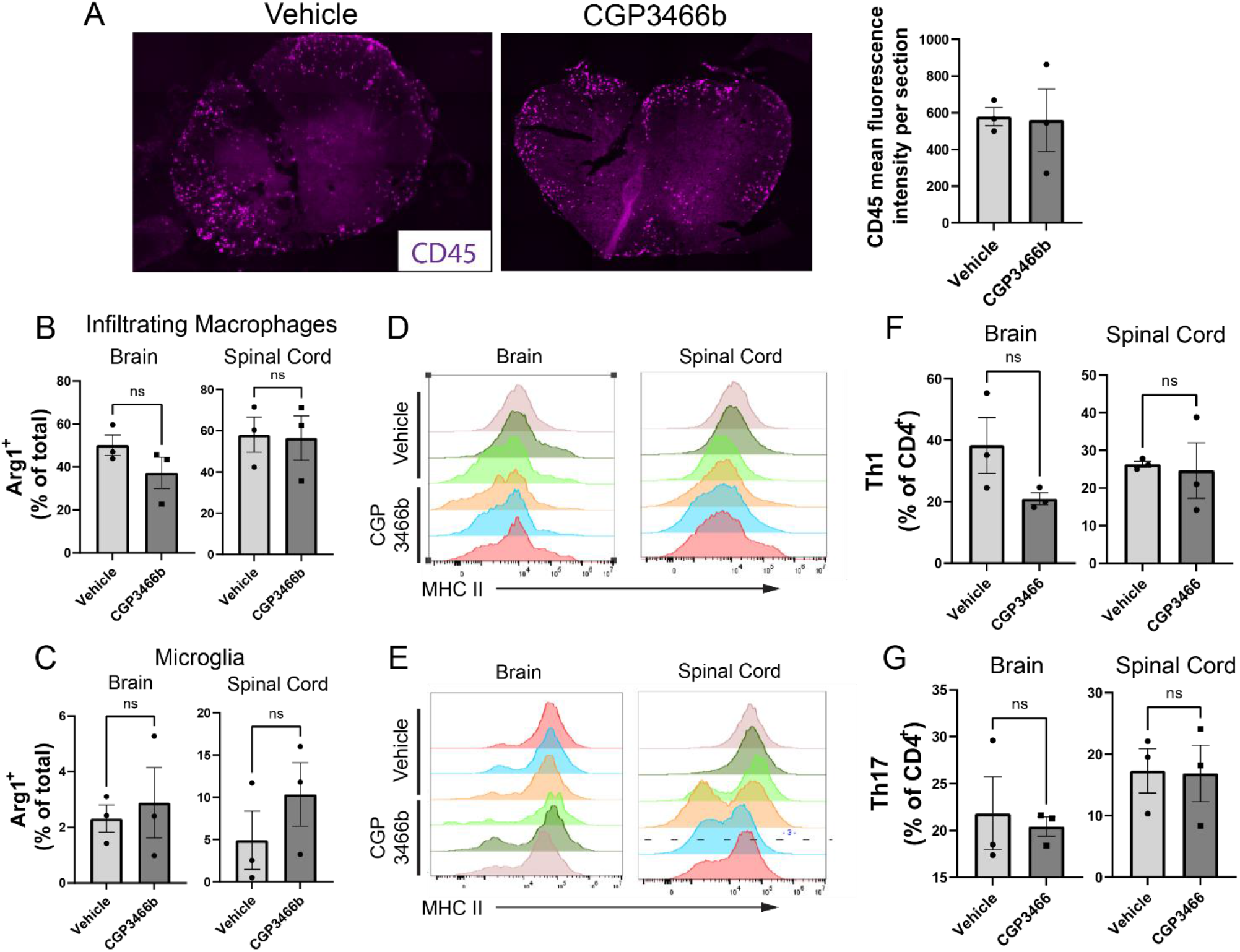
CGP3466b does not impact CNS immune infiltration in MOG_35-55_ EAE. **(A)** CD45+ infiltrates in lumbar spinal cord at post-immunization day 19. Mice were treated with vehicle or 4 mg/kg CGP3466b daily starting on post-immunization day 0. A representative image is shown (*left*), along with quantification performed from n=3 mice per group (*right*). **(B)** Infiltrating macrophages and **(C)** resident microglia in the brain and spinal cord were examined via flow cytometry for expression of arginase-1 as a percentage of total macrophages or microglia, respectively. Data represent mean ± SEM of 3 mice per group. **(D)** Infiltrating macrophages and **(E)** resident microglia were examined via flow cytometry for expression of MHC II, shown as mean fluorescence intensity. Data shown from 3 mice per group. **(F)** Th1 cells and **(G)** Th17 cells were quantified via flow cytometry in the brain and spinal cord as a percentage of total CD4+ cells. Data represent mean ± SEM of 3 mice per group. All statistical analyses performed with student’s t-test.

## 3 Discussion

In this study, we evaluated the GAPDH nitrosylation inhibitor CGP3466b as a potential novel therapy for MS. We found that in the MOG_35-55_ EAE model of MS, GAPDH is a major target of nitrosylation within the CNS, which is selectively blocked by systemic treatment with CGP3466b. We further showed that CGP3466b is neuroprotective in the context of neuroinflammation, attenuating disease course in our murine model of MS and reducing axonal damage in the optic nerve. Finally, we showed that CGP3466b has minimal effects on the immune response *in vitro* and *in vivo*, suggesting that CGP3466b is directly neuroprotective and acts independently of the peripheral immune system.

Modest effects of CGP3466b were observed on myeloid cell function only at a high dose (200 nM). Although CGP3466b blocks GAPDH nitrosylation at low nanomolar concentrations (Hara et al., 2006), at higher concentrations the drug has other targets such as the PCMT1/MST1 signaling pathway (Liang et al., 2017) and succinate-dependent H_2_O_2_ release from mitochondrial complex I (Zoccarato et al., 2008). These lower-affinity targets may explain the observed effects on myeloid cells at high dose. However, it is unlikely that such a high concentration is achieved *in vivo* with the 4 mg/kg dose used in our studies.

There has been longstanding interest in blocking NO-mediated pathways as a means to protect oligodendrocytes and neurons in MS. Microglia, CNS-resident macrophages, and astrocytes – the primary sources of NO – represent a common link in the pathology of relapsing and progressive MS. As previously noted, NO levels increase in active MS lesions and during relapse, leading some to investigate NO as a biomarker (abdel Naseer et al., 2020). Finally, NO directly contributes to neuro-axonal injury in animal models of neuroinflammation (Nikic et al., 2011). Nonetheless, NO has pleiotropic effects in inflammation, some of which are required for resolution of inflammation or simply for normal function of the nervous system, such that limiting NO production altogether has yielded inconsistent results (Encinas et al., 2005). In contrast, selectively targeting SNO-GAPDH with CGP3466b blocks a specific NO-mediated signaling pathway involved in neuroaxonal injury while preserving other functions of NO.

SNO-GAPDH has been shown to trans-nitrosylate several proteins, including the deacetylating enzyme sirtuin-1 (SIRT1) (Kornberg et al., 2010). SIRT1 is a key regulator of mitochondrial biogenesis (Chuang et al., 2019; Majeed et al., 2021), and SNO-GAPDH has also been shown to trans-nitrosylate several mitochondrial enzymes (Kohr et al., 2014). Therefore, it is possible that blocking CGP3466b is protective in neurons and axons because it prevents SNO-GAPDH-mediated mitochondrial dysfunction. Future studies will examine the effect of CGP3466b on mitochondria health in neurons, along with other potential mechanisms underlying its neuroprotection.

Because CGP3466b has been evaluated in Phase II clinical trials, there is an established safety profile and low threshold for clinical translation. Although CGP3466b did not have a significant effect on the clinical course of Parkinson’s disease in humans, this is likely because Parkinson’s disease pathology is complex with much less dependence on oxidative/nitrosative stress than the MPTP mouse model in which CGP3466b showed benefit. In contrast, MS is a primary inflammatory disease in which macrophage/microglia and astrocyte generation of NO is a known aspect of pathology in both the human disease and the EAE mouse model. As such, the rationale for involvement of SNO-GAPDH in human disease is much stronger in the case of MS.

All disease modifying therapies currently approved for MS act by targeting the immune system – primarily by limiting peripheral immune activation or preventing peripheral immune cells from infiltrating the CNS (Callegari et al., 2021). However, none of these medications limit CNS damage during a relapse or, most importantly, slow disability accrual in non-active progressive MS, largely because progressive MS pathology is driven by compartmentalized innate immune activation instead of the T-cell driven attack seen in relapsing-remitting MS (Lassmann et al., 2012; Rice et al., 2013). Therefore, neuroprotective therapies that prevent inflammatory CNS injury represent a major goal in MS research, regardless of the source or stage of inflammation. Such therapies might be used as adjunct treatments in combination with immunomodulatory therapies in relapsing MS and/or as first-in-class agents to slow neurodegeneration and therefore disability accrual in progressive MS. Future studies will evaluate CGP3466b as an adjunct therapy in murine models of MS.

In summary, our data indicate that blocking GAPDH nitrosylation with CGP3466b significantly attenuates disability in the murine EAE model of MS through a neuroprotective mechanism independent of the peripheral immune system. Given the substantial need for neuroprotective agents as adjunct therapies in both relapsing and progressive MS and due to its established safety profile, CGP3466b holds promise as a therapeutic strategy in MS that merits further investigation.

### 3.1 Materials and Methods

#### Mice

Wild-type C57BL/6J mice were purchased from the Jackson Laboratory (stock # 000664) and housed in a dedicated Johns Hopkins rodent facility with regulated temperature (20-22°C), 50% humidity, and a 12-hour light/12-hour dark cycle with free access to water and solid food. Protocols were approved by the Johns Hopkins Institutional Animal Care and Use Committee (Protocol MO21M370).

#### Antibodies

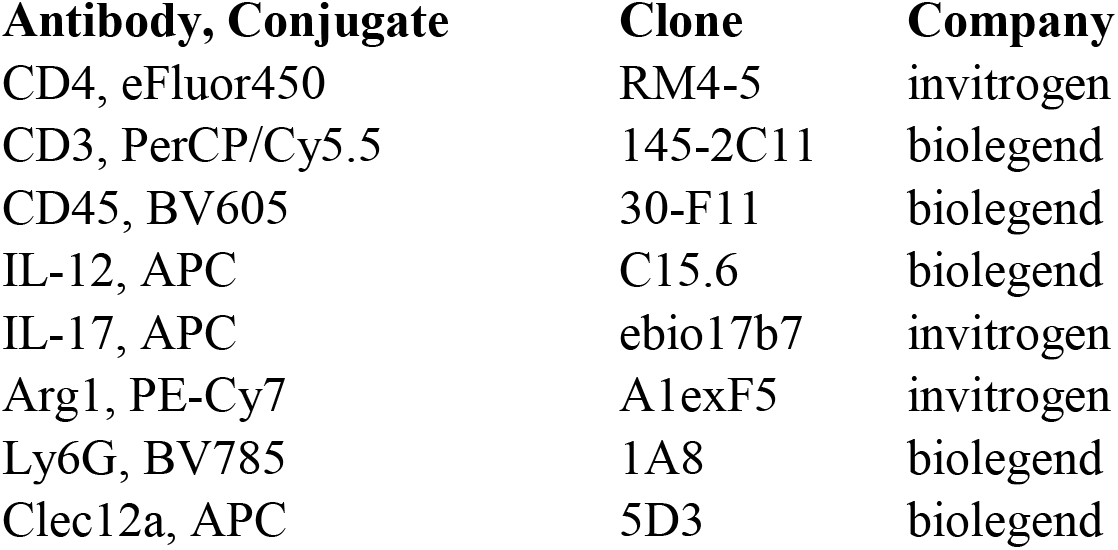
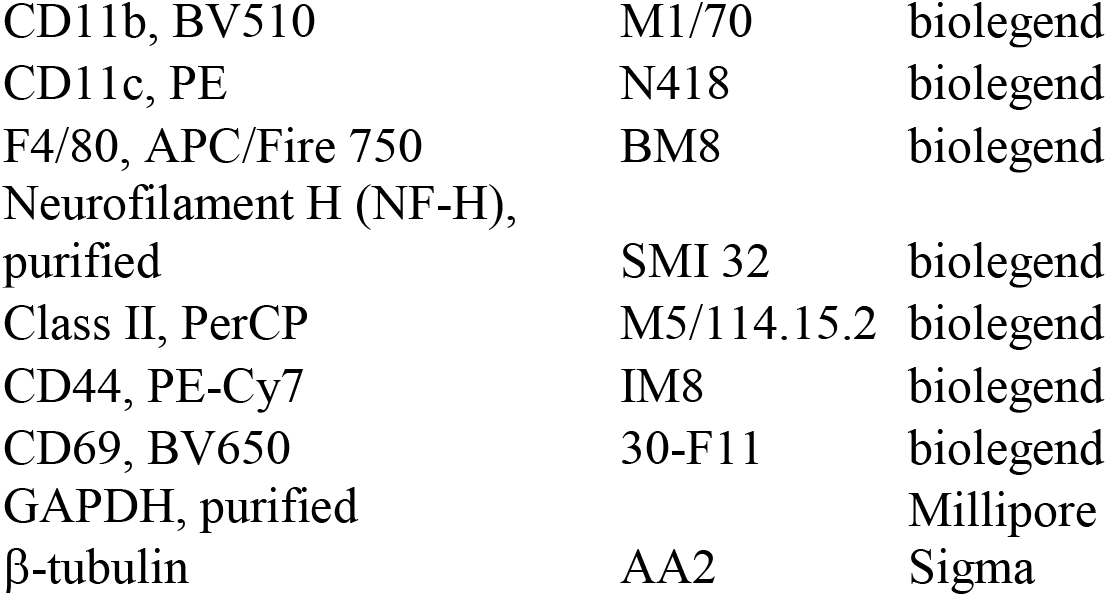

#### EAE induction and scoring

Active EAE was induced in female C57BL/6J mice (8–12-week-old) after 1-week acclimatization to the animal facility. MOG_35–55_ peptide dissolved in PBS at a concentration of 2 mg/mL was mixed 1:1 with complete Freund’s adjuvant (8 mg/ml Tuberculin toxin in incomplete Freund’s adjuvant), then mixed for 10 minutes into an emulsion. On day 0, mice were immunized by injecting 50 μl of the emulsion subcutaneously into each of two sites on the lateral abdomen. In addition, on day 0 and again on day 2, mice were injected intraperitoneally with 250 ng pertussis toxin. Control animals were treated with complete Freund’s adjuvant and pertussis toxin alone. Scoring was performed in a blinded manner according to the following scale: 0, no clinical deficit; 0.5, partial loss of tail tone; 1.0, complete tail paralysis or both partial loss of tail tone plus awkward gait; 1.5, complete tail paralysis and awkward gait; 2.0, tail paralysis with hind limb weakness evidenced by foot dropping between bars of cage lid while walking; 2.5, hind limb paralysis with little to no weight-bearing on hind limbs (dragging), but with some movement possible in legs; 3.0, complete hind limb paralysis with no movement in lower limbs; 3.5, hind limb paralysis with some weakness in forelimbs; 4.0, complete tetraplegia but with some movement of head; 4.5, moribund; and 5.0, dead.

#### Treatment of mice with CGP3466b

CGP3466b maleate (Tocris, cat # 2966) stock solution (50 mM in DMSO) was diluted to 4% DMSO in PBS as a working solution (final CGP3466b concentration 20 - 800 µg/ml depending on dosage to be delivered). Mice were treated daily with i.p. injection of drug or an equal volume of vehicle control (4% DMSO in PBS). Treatment either began on the same day as immunization with MOG_35–55_ (PID 0) or on PID 10.

#### Biotin switch assay

The assay was performed as previously described with slight modifications (Jaffrey & Snyder, 2001). Briefly, spinal cords were flushed from the spinal column with hydrostatic pressure and then then dissolved in RIPA/HEN (HEN buffer adjusted to contain 1% Triton X-100, 1% sodium deoxycholate, 0.1% SDS) with a hand-held tissue homogenizer. The homogenate was clarified by spinning at 13,000g for 10 min at 4°C. Supernatant was collected and stored at -80°C until use. Free thiol groups were blocked with 20 mM MMTS (methyl methanethiosulfonate) for 20 min. Protein was precipitated in acetone and then resuspended in a labeling solution containing 50 mM sodium ascorbate and 0.8 mM biotin-HPDP for 1 hour. Protein was precipitated in ice-cold acetone, resuspended, and pulled-down overnight with high-capacity neutravidin agarose beads. The beads were washed and biotinylated proteins were eluted with HEN/10 buffer (HEN buffer diluted 1/10 in water) containing 1% beta-mercaptoethanol (Sigma). Protein was resolved by SDS/PAGE. Bands were transferred to PVDF Immobilon P membranes using a wet transfer, blocked in TBS-T containing 5% milk, and probed overnight at 4°C with primary antibodies. Blots were then washed and stained with HRP-conjugated secondary antibody (Jackson ImmunoResearch). Immunoblots were visualized using the SuperSignal West ECL system (ThermoFisher) followed by film exposure. Bands were quantified with ImageJ software.

#### Immunofluorescence imaging

Mice were euthanized by overdose of isoflurane (adjusting the isoflurane flow rate to 5% until breathing stopped) and then perfused via cardiac puncture with ice-cold PBS followed by a paraformaldehyde perfusion. After perfusion, optic nerves and spinal cords were dissected out. Tissue was fixed in paraformaldehyde overnight, sucrose protected with 30% sucrose, and frozen in OCT. Cryosections (12 µm) from proximal optic nerve were incubated with anti-SMI-32 antibodies overnight, washed with PBS-T, and incubated with anti-mouse secondary antibody (AlexaFluor 647). Spinal cord sections (12 µm) were incubated with anti-CD45 (APC conjugated) antibody overnight at 4°C. Sections were imaged on a Zeiss Axio Observer Z1 epifluorescence microscope and Zeiss 710 confocal microscope with the appropriate excitation and emission filters. SMI-32 staining was quantified with ImageJ software.

#### Isolation, culture, and treatment of murine bone marrow derived dendritic cells (BMDCs) and bone marrow derived macrophages (BMMs)

Cells were isolated as described (Lutz et al., 1999). Briefly, femurs were removed from 6–10-wk-old C57BL/6J mice, cut on both ends, and marrow was flushed with PBS. Bone marrow cells were then pelleted at 1500 rpm, resuspended in red blood cell lysis buffer, and after 1 minute the reaction was quenched with excess PBS. After another centrifugation, cells were resuspended in complete RPMI media consisting of RPMI-1640 with GlutaMAX supplement, along with 10% FBS, 1% penicillin-streptomycin, and 50 μM beta-mercaptoethanol (Sigma). On day 0, ∼2 × 10^6^ cells were then seeded per 100-mm plate in 10 mL media containing 20 ng/mL recombinant murine M-CSF (Peprotech). On day 3, an additional 10 mL fresh cRPMI media containing 20 ng/mL M-CSF was added to each plate. On day 5, half the culture supernatant from each plate was removed and centrifuged, and the pelleted cells resuspended in 10 mL fresh cRPMI with 20 ng/mL M-CSF and added back to the plates. On day 8, all nonadherent cells (representing the BMDC fraction) were collected, pelleted by centrifugation, resuspended in fresh cRPMI media with 20 ng/mL GM-CSF, and plated into 96-well dishes. These BMDCs were then treated overnight with or without LPS (Sigma) 100 ng/ml plus the indicated doses of CGP3466b (dissolved in DMSO) or vehicle (DMSO alone). The following day (day 9), these cells were then assayed by flow cytometry. Adherent macrophages (BMMs) were removed from the petri dish with macrophage detachment solution (Promocell) and replated into 6 well plates along with LPS 100 ng/mL and the indicated doses of CGP3466b or vehicle. These cells were then assayed by flow cytometry or cytokine ELISA.

#### Preparation of tissue for flow cytometry

Mice were euthanized by overdose of isoflurane (adjusting the isoflurane flow rate to 5% until breathing stopped) then perfused with ice-cold PBS via cardiac puncture. Brain and spinal cord were collected, mechanically dissociated, then chemically dissociated with collagenase and DNase with constant shaking. Cells were then passed through a 100 µM filter and myelin debris was removed by resuspending the cell pellet with a debris removal solution (Miltenyi Biotec), overlaying with PBS, and spinning at 3000g, then removing the myelin debris layer. Cell pellets were then resuspended in PBS and stained with antibody as described below.

#### CD4+ T-cell isolation and culture

T-cells were isolated by generating single-cell suspensions from the spleen and lymph nodes of 6-10-week-old C57BL/6 mice. Spleens were disrupted with the plunger of a syringe over a 70-μm nylon cell strainer (BD Falcon), then cells were pelleted by centrifugation (500 × g for 5 min) and resuspended in fresh FACS buffer. CD4+ cells were then isolated by negative selection using the Mojo CD4+ cell isolation kit from Biolegend (Catalog 48006) according to the manufacturer’s instructions. Plates were coated overnight at 4°C with anti-mouse CD3 antibody (3 µg/mL). T cells were resuspended in complete RPMI (RPMI plus 10%FBS, 1% Pen-strep, 1% glutamax, 0.1% 2-mercaptoethanol) along with 3 ug/mL anti-mouse CD28 antibody. Cells were cultured for 96 hours and treated for the last 24 hours with vehicle or CGP3466b before proceeding with flow cytometry staining.

#### ELISA

After overnight stimulation with LPS 100 ng/mL ± CGP3466b or vehicle, BMM culture supernatants were collected and cytokine production was assayed using ELISA kits for IL-1β, IL-6, and IL-10 purchased from eBioscience, according to manufacturer’s instructions. Plates were read at 450 nm on a tabletop colorimetric spectrophotometer.

#### Flow cytometry staining

For panels involving T cells, cells were first resuspended in a solution of complete RPMI with brefeldin/Monensin and PMA/ionomycin. BMDCs and BMMs were treated with brefeldin and monensin. These treatments lasted 4 hours at 37°C.

All staining was performed in the dark at room temperature. Cells were first stained with zombie NIR (1:1000) for 10 minutes along with CD16/32 (1:200) dissolved in PBS. Cells were then washed and resuspended in a solution of PBS+2% FBS+1 mm EDTA and stained with the relevant antibodies. All surface antibodies were stained at a 1:300 dilution. Cells were then washed and incubated in FoxP3 fixation/permeabilization buffer following the manufacturer recommended protocol. Intracellular staining was performed with conjugated antibodies (1:200) against the specified proteins in permeabilization buffer for 1 h, washed twice, and then analyzed cells with a Cytek Aurora Flow cytometer. Data analysis was performed with FlowJo software.

#### Statistical Analyses

All statistical analyses were performed using GraphPad Prism software. Details of statistical analyses for each experiment can be found in the figures and figure legends.

## 5 Author Contributions

WHG contributed to project and experimental design, data acquisition, data analysis/interpretation, and drafting of the manuscript and figures. SH, PG, and EA contributed to data acquisition. MDK contributed to project and experimental design, data acquisition, data analysis/interpretation, drafting/editing of the manuscript and figures, and provided funding support.

## 7 Acknowledgments

The authors would like to thank Matthew Smith, Bindu Paul, Judy Lee, Michelle Taylor, Thomas Garton, and Em Harrington for technical assistance.

