## Supplementary Figure 1 for "Therapeutic Potential of Blocking GAPDH Nitrosylation with CGP3466b in Experimental Autoimmune Encephalomyelitis"

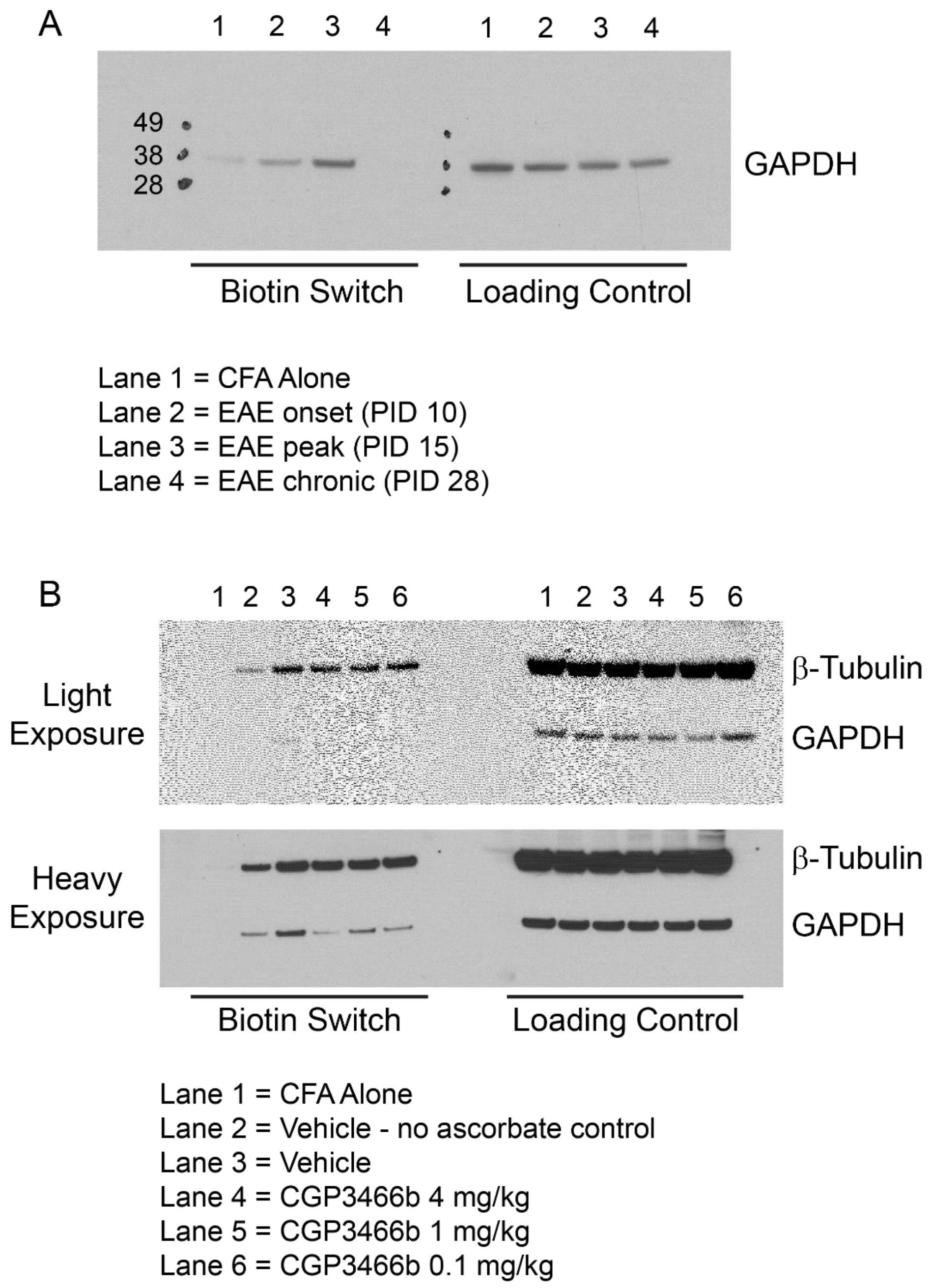


**Supplementary Figure 1.** Full immunoblots from Figure 1. **(A)** Full blot from Figure 1A. **(B)** Full blot from Figure 1B. For (B), biotin switch and loading control samples were run on different gels but exposed on same film, and blots were cut between molecular weight markers 38 and 49 kD to probe for GAPDH and β-Tubulin, respectively.
